# Distinct roles for partially redundant transcription factors in *Caenorhabditis elegans* mesoderm lineage development

**DOI:** 10.64898/2026.09.01.748736

**Authors:** Yuntian Gan, Erin L. Marble, Eli Preston, Brenna Tento, Emily Jiang, John Isaac Murray, Priya Sivaramakrishnan

## Abstract

Developmental transcription factors often have overlapping functions, making it difficult to define the distinct roles of individual factors during lineage specification. We investigated the partially redundant transcription factors TBX-35 and CEH-51 in the *Caenorhabditis elegans* embryonic MS mesodermal lineage using 4D lineage tracing, reporter imaging, genetics, and single-cell RNA sequencing. In *tbx-35* mutants, MS descendants showed progressively slower cell cycles and a division pattern that increasingly resembled the cousin C lineage. Fate-regulator expression also shifted toward C-like features, including ectopic *pal-1* and expanded HLH-1 expression, although mutant cells did not simply adopt normal C-lineage positions. Loss of *tbx-35* also impaired a later MS-dependent Notch induction in the AB lineage while leaving an earlier induction intact. CEH-51 showed a different pattern of activity whereby its protein became enriched in anterior MS daughters, and *ceh-51* mutants produced later, more restricted lineage defects that were strongest in descendants of cells with higher CEH-51 levels. Single-cell profiling identified overlapping but nonidentical sets of genes dependent on the two factors. TBX-35-dependent changes were strongest at earlier stages, whereas CEH-51-dependent genes became more prominent later and were enriched in anterior MS sublineages. Finally, temperature-shift experiments determined that the severity and onset of *tbx-35* mutant phenotypes depend on the maternal temperature environment and cannot be explained by differences in residual CEH-51 expression. These findings reveal that TBX-35 and CEH-51 contribute differently across the MS lineage and that reliable mesoderm development is supported by overlapping zygotic and maternal regulatory inputs.

## Introduction

Animal development relies on transcription factors that are reused across tissues and developmental stages and often act in partially redundant combinations. Such redundancy can make development robust to perturbation, but it also obscures regulatory logic. Two central questions are whether partially redundant transcription factors have shared or distinct targets, and how their functions are partitioned between cell-fate specification and other dynamic properties of a developing lineage, including division timing, cell death, and cell positioning.

The invariant cell lineage of the *Caenorhabditis elegans* embryo provides an unusually direct system in which to address these questions. Every embryonic cell can be identified by its division history (“lineage”), and phenotypes such as transcriptional state, reporter expression, division timing, and cell position can all be mapped onto the lineage tree^1^. Exhaustive single-cell transcriptomic atlases^2–5^, lineage-resolved reporter resources^6–8^, and imaging-based 4D lineage tracing^9–12^ make it possible to connect regulatory perturbations to both molecular and developmental phenotypes at single-cell resolution.

Here, we dissect the roles of the partially redundant transcription factors TBX-35/T-box and CEH-51/NK-2- like in the *C. elegans* embryonic mesodermal MS lineage. This system allows us to determine how these factors partition their regulatory functions across developmental time and cell type and relate this to cell fate specification and other lineage phenotypes such as division timing. The MS lineage generates pharyngeal and body-wall muscle, coelomocytes, GLR glia, and other mesodermal cell types^1^. MS identity is established by a compact transcriptional cascade (Fig. 1A)^13^. The gene regulatory network that specifies the MS blastomere begins with the maternal factor SKN-1 promoting transcription of the redundant GATA factors MED-1 and MED-2 in EMS (the parent of MS and the endodermal lineage E). These in turn activate the T-box factor TBX-35 in the MS cell at the 8-cell stage. TBX-35 and other factors then activate CEH-51 expression in the MS daughters^14–17^.

**Figure 1.**
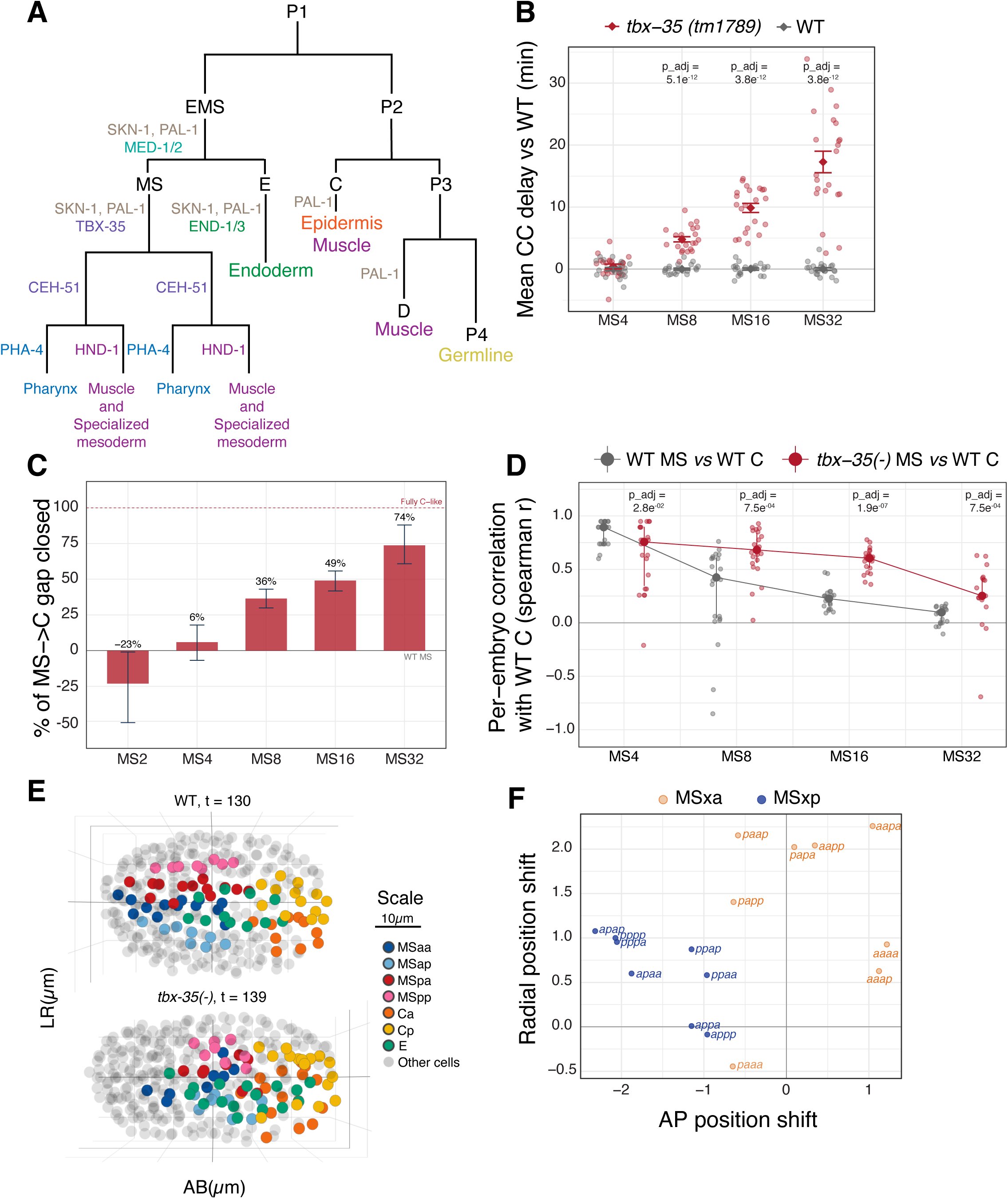
Loss of *tbx-35* shifts the MS lineage toward a C-like division program. (A) Schematic of the *C. elegans* founder lineage, with broad fates and the key transcription factors specifying each MS branch indicated. (B) Mean cell-cycle delay of MS-lineage divisions in *tbx-35(-)* (red) relative to the wild-type mean (grey) at MS4, MS8, MS16 and MS32. Each point is one embryo’s mean, diamond and bars are mean ± SE. n = 22 *tbx-35(-)*, n = 22 wild type. p values from Wilcoxon rank-sum, BH-corrected (MS8/16/32 shown; MS4 is n.s.). (C) Fraction of the MS-to-C cell-cycle-timing gap closed by *tbx-35(-)* MS cells at each stage (0% = wild-type MS timing, 100% = homologous wild-type C timing). Bars are mean, whiskers are bootstrap 95% CI (2000 resamples of embryos, wild type and mutant independently). n = 22 *tbx-35(-)*, n = 22 wild type. (D) Per-embryo rank correlation (Spearman r) between MS cell-cycle lengths and their lineage-homologous wild-type C cells, for wild-type MS (grey) and *tbx-35(-)* MS (red). Diamond and bars are median ± IQR. n = 17-22 *tbx-35(-)* per stage, n = 22 wild type per stage. p values from Wilcoxon rank-sum (mutant vs wild type), BH-corrected. (E) Aligned 3D renderings of MS- lineage cell positions in one wild-type (t = 130) and one severe *tbx-35(-)* (t = 139) embryo, approximately stage-matched, colored by MS4 sublineage (MSaa, MSap, MSpa, MSpp), E, and the C sublineages (Ca, Cp); other cells grey. AB, anterior-posterior; LR, left-right. Scale bar, 10 um. Representative single embryos. (F) Displacement of MSxa (orange) and MSxp (blue) cells in *tbx-35(-)* relative to wild type, radial (surface- interior) vs anterior-posterior shift; each point is one cell (mean shift across embryos, position at birth). n = 22 *tbx-35(-)*, compared to the wild-type reference (Richards et al. 2013).

In the absence of *tbx-35*, the MS lineage fails to produce some MS-derived cell types, and a subset of embryos derepresses PAL-1/Caudal, a regulator of the C lineage (a neighboring early embryonic lineage that normally produces posterior body-wall muscle and hypodermis)^18,19^. However, some MS-derived cell types such as coelomocytes are still produced in *tbx-35(-)* embryos, suggesting a partial or incompletely penetrant transformation of MS to a C-like fate^17^. *ceh-51* mutants have milder defects, with many MS-derived tissues specified but abnormalities in pharynx morphology, muscle positioning, and coelomocyte number. *tbx-35; ceh-51* double mutants have a stronger phenotype than *tbx-35* alone and phenocopy the *med-1;med-2* double mutant^16^.

These observations raise several questions about how TBX-35 and CEH-51 orchestrate MS development. Prior studies identified several early regulatory defects, but much of the phenotypic characterization focused on terminal cell types, leaving unclear how these defects unfold across the developing MS lineage. If *tbx-35* mutant MS cells partially acquire C identity, do they also acquire C-like division behavior? Does CEH- 51 simply reinforce TBX-35 activity, or does it make distinct temporal or sublineage-specific contributions? More broadly, it remains unknown whether the partial redundancy of TBX-35 and CEH-51 reflects regulation of shared downstream genes or overlapping control of distinct developmental programs.

Here, we use complementary lineage, genetic, and transcriptomic approaches to define how TBX-35 and CEH-51 shape MS development. By relating lineage phenotypes to transcription-factor expression and mutant transcriptional profiles, we determine how these partially redundant regulators contribute to lineage behavior, cell-fate specification, and developmental robustness.

## Results

### Loss of *tbx-35* shifts the MS lineage toward a C-like division program

Prior work with cell-fate reporters indicated that *tbx-35* null mutants and *tbx-35*; *ceh-51* double mutant MS lineages are partially transformed toward a C-like fate, including ectopic zygotic PAL-1 activation, ectopic hypodermal cells and mispositioned PAL-1-dependent body-wall muscle^16,17^. How this transformation extends to the dynamic behavior of each cell in the lineage - their division timing, division pattern, and cell positions, is unknown. For this analysis, we collected time-lapse 4D images of *tbx-35(tm1789)* null mutant (hereafter referred to as *tbx-35(-)*) embryos expressing mCherry::histone and performed 4D lineage tracing with StarryNite and AceTree (^9,11^, Fig. S1A). Because wild-type C descendants divide on average ∼40% more slowly than MS descendants, a transformation of MS to a C-like identity predicts slower MS divisions^20^. Indeed, MS descendants divided progressively later in mutants, slowing by on average 5 min (17%) at MS8, 10 min (23%) at MS16, and 17 min (29%) at MS32 (Fig. 1B). The delay was mostly restricted to the MS lineage, although we observe a significant delay in the sister lineage E at the E8 stage (11% of the wild-type cell cycle, Fig. S1B). As *tbx-35* is not expressed in E, this is likely a nonautonomous defect, potentially reflecting the concurrent mispositioning of the E lineage by its defective MS sister.

To quantify how far the mutant lineage shifts towards C-like division rates, we expressed each stage’s mean cell-cycle length on a scale running from wild-type MS timing (0%) to that of the homologous wild-type C cell (100%), i.e., the fraction of the MS-to-C timing gap that is closed. By this measure the mutant MS lineage closed 36% of the gap at MS8, rising to 74% by MS32 (Fig. 1C). Thus, the shift toward C-like division rate deepens over developmental time. The transformation to C is also reflected in division pattern, not only division timing. Wild-type MS and C lineages have distinct division architectures. MS is largely synchronous with a mild posterior-slower gradient through MS16 and pronounced asynchrony at MS32/MS64, whereas in C the myogenic Cxp branches divide more slowly than the ectodermal Cxa branches and also divide one extra time (Fig. S1A)^1,19,20^. To ask whether mutant MS adopts a C-like pattern of fast and slow cells independent of overall rate, we correlated each MS cell’s cell-cycle length with that of its corresponding C cell across embryos. In wild-type embryos, this correlation is weak and decreases across time, but in *tbx-35(-)* mutants MS division times become more correlated with those of the homologous C cells (Fig. 1D, S1C). This emerges by MS8 and is clearest at MS16 (Spearman r = 0.60 in mutants compared with 0.23 in wild-type, Fig. 1D). The strongest example is Caapa, a C-lineage cell that divides more than an hour after its cousins in wild type. The corresponding MS cell, MSaapa, switches from one of the fastest wild-type MS16 divisions to the slowest in *tbx-35* mutants, which represents a 33-min average delay in mutants, compared to an average ∼7 min delay for the other MS16 cells. The magnitude of delay differs widely across embryos (12–76 min), consistent with a variably penetrant adoption of Caapa- like behavior (Fig. S2A). Both the global cell-cycle delays and the correlation with C lineage division patterns were reduced but not eliminated in *tbx-35(+)* array rescued embryos (Fig. S2B, S2C).

We next asked whether MS-derived cells adopt C-like cell positions in *tbx-35* mutant embryos (Fig. 1E,F). We first measured position defects based on each cell’s distance from its normal position along the anterior-posterior and radial axes, as well as displacement from its wild-type neighbors using a nearest-neighbor deviation (NN) score. The *tbx-35(-)* mutant MSxa-derived cells remain closer to the surface than their more central location in wild-type embryos, suggesting a defect in their internalization during gastrulation (Fig. 1F). They also had relatively subtle anterior-posterior position defects (Fig. 1E,F). The corresponding Cxa cells produce epidermal fates and remain on the surface in both genotypes. Mutant MSxp-derived cells are mispositioned anteriorly, with more moderate surface shifts, consistent with a previous finding of accumulation of muscle cells in the middle of the embryo^17^ (Fig. 1E).

Both epidermal (Cxa) and muscle (Cxp) cells from the C lineage remain further posterior than the corresponding *tbx-35(-)* MS-derived cells, possibly due to incomplete fate conversion or birth-position effects. Similarly, the space normally occupied by the MS lineage cells instead contained cells derived from other lineages such as ABa and ABp, suggesting that the mispositioning of MS-derived cells displaces cells from other lineages. Throughout the MS lineage, cells became more mispositioned relative to their wild-type neighbors at later stages, but this mispositioning was already significant by MS4, indicating regulation of cell positioning by lineage identity early in MS development (Fig. S2D). Together, *tbx-35* mutant MS cells acquire more C-like division timing, and while their positions have some C-like characteristics, they do not simply map onto the normal positions of corresponding C-lineage descendants.

### CEH-51 expression shows anterior bias that is lost in *tbx-35(-)* mutants

Because residual CEH-51 activity was proposed to explain the partial penetrance of tbx-35 mutants, we next quantified endogenous CEH-51 protein dynamics in wild-type and *tbx-35(-)* embryos by StarryNite lineage tracing of an endogenous CEH-51::GFP CRISPR reporter. Consistent with prior observations^8,21^, we observed expression throughout the MS lineage starting in the MS granddaughters (MS4) and at high levels in all MS8 and MS16 cells (Fig. 2A,B). We noticed a strong quantitative bias within the MS lineage, with the anterior daughter expressing more CEH-51::GFP than the posterior daughter after many divisions. This asymmetry builds at MS4, is strongest at MS8–MS16 and is weaker by MS32 (Fig. 2A-C, S3B). The anterior daughter shows substantially more nuclear CEH-51 within a few minutes of the division, before newly synthesized GFP could mature, suggesting that the asymmetry is controlled at least partly post-transcriptionally (Fig. 2C). The asymmetry persists longest in the anterior daughters of the terminal divisions (MSaaaa, MSpaaa, and MSxppa), where CEH-51::GFP remains detectable past the 350-cell stage, whereas their posterior sisters (MSaaap, MSpaap, MSxppp) lose it by the end of the 200-cell stage (Fig. S3C).

**Figure 2.**
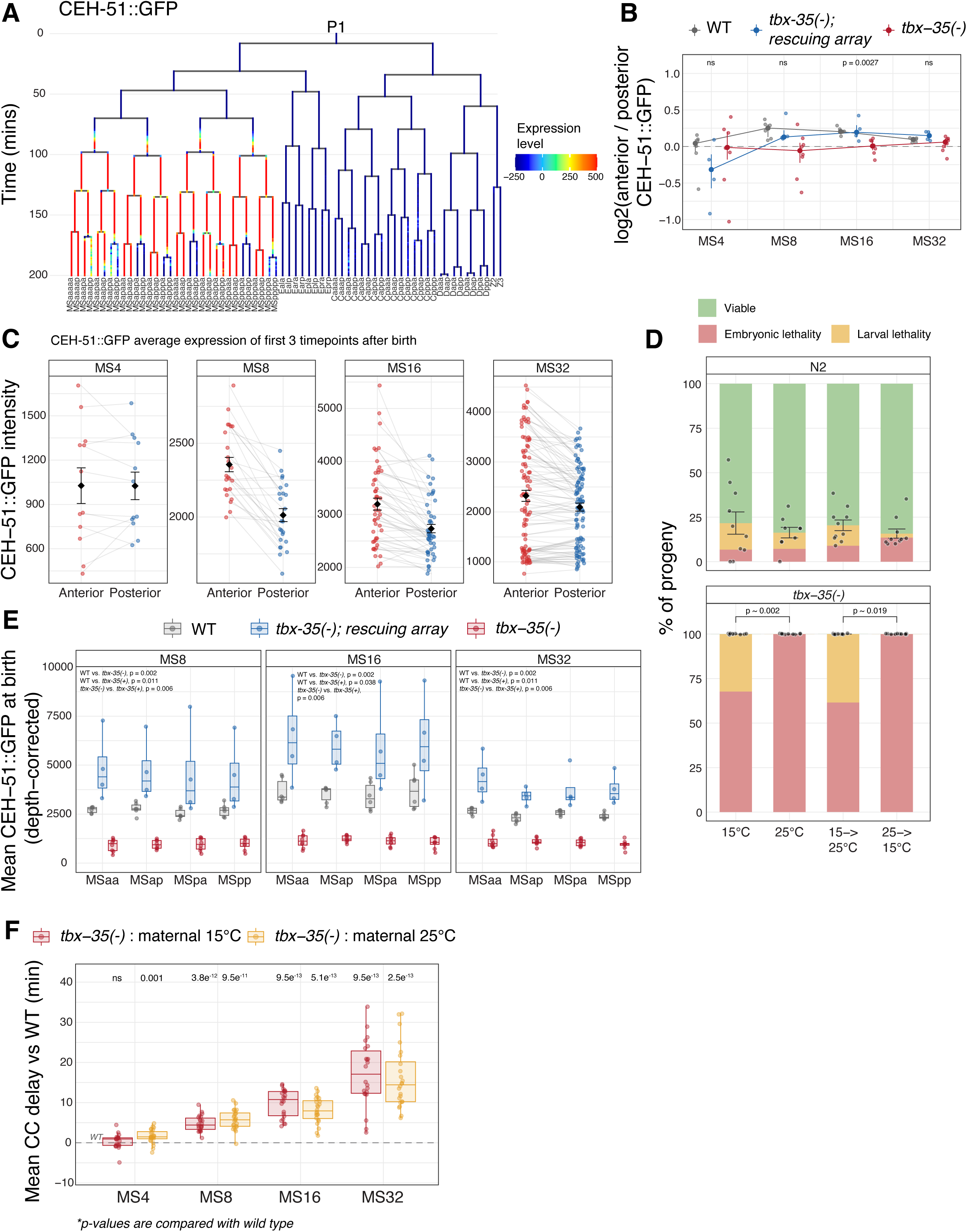
Asymmetric CEH-51 expression and temperature-dependent lethality in *tbx-35(-)*. (A) CEH-51::GFP expression tree for a representative wild-type embryo (P1 root); branch color, depth- corrected CEH-51::GFP; y-axis, time. (B) Per-embryo mean log2(anterior/posterior CEH-51::GFP) per stage, in wild type (grey), *tbx-35*(-); rescuing array (blue), and *tbx-35(-)* (red). Diamond and bars are median ± IQR; dashed line at zero is symmetry. n = 6 wild type, n = 4 *tbx-35*(-); rescuing array, n = 8 *tbx-35(-)*. p values from Wilcoxon rank-sum. (C) Anterior vs posterior CEH-51::GFP intensity (depth-corrected, mean of the first 3 timepoints after birth) for every wild-type MS division at MS4-MS32. Grey lines connect the two daughters of one division (anterior red, posterior blue); black, mean ± SE. n = 6 wild type. (D) Combined lethality. Stacked bars are the fraction of progeny that were viable (green), larval-lethal (amber), or embryonic-lethal (red), for N2 and *tbx-35(-)* at constant 15°C or 25°C or shifted at the 2-4-cell stage (15->25, 25->15). Points, individual broods; error bar, SE of total lethality. n = 9-10 broods per condition. p values from Wilcoxon rank-sum (total lethality). (E) Mean CEH-51::GFP at birth (depth-corrected) per MS4 sublineage at MS8/MS16/MS32, for wild type (grey), *tbx-35*(-); rescuing array (blue), and *tbx-35(-)* (red). Boxes are median/IQR, points are per-embryo. n = 6 wild type, n = 4 *tbx-35*(-); rescuing array, n = 8 *tbx- 35(-)*. p values from Wilcoxon rank-sum, BH-corrected, comparing each embryo’s mean over all sublineages at that stage. (F) Mean cell-cycle delay vs wild type for *tbx-35(-)* MS divisions by stage, from mothers reared at 15°C (red) and 25°C (amber). Boxes are per-embryo means (median/IQR); dashed line, wild type (delay = 0). n = 22 maternal 15°C, n = 24 maternal 25°C. p values compare each temperature to wild type (Wilcoxon rank-sum, BH-corrected); 15°C and 25°C did not differ at any stage.

CEH-51::GFP is reduced by variable amounts across *tbx-35(-)* embryos (Fig. 2E, S3A). Residual expression was detectable in most imaged embryos (14/16), with only 2 embryos showing near complete absence of CEH-51 (Fig. S3D,E). When present, expression was stronger in the anterior MSa-derived lineages (Fig. S3D). This result is consistent with Broitman-Maduro et al., 2009, who reported some remaining CEH-51 reporter expression in *tbx-35(-)* that was reduced and lower, with onset undetectable in 52% of embryos. The *tbx-35*-independent CEH-51 expression could contribute to the milder phenotype of *tbx-35* single mutants relative to *ceh-51*; *tbx-35* double mutants^16^. However, loss of *tbx-35* reduces or eliminates CEH-51 expression asymmetry. Across all divisions, CEH-51::GFP in *tbx-35(-)* shows no bias for anterior daughters at any stage and is significantly different from wild-type expression at MS16, when the wild-type asymmetry is strongest (Fig. 2B). The rescuing *tbx-35(+)* array partially restores the asymmetry (Fig. 2B).

### Temperature impacts *tbx-35(-)* embryonic lineage phenotypes

Past work found that *tbx-35(-)* embryos have stronger phenotypes at higher temperatures, with most embryos arresting after elongation at 15°C, but prior to elongation at 20°C. Also, fewer *tbx-35(-)* cells expressed MS-specific cell type markers at 23°C, suggesting a more severe phenotype at higher temperatures^16^. We confirmed and extended this effect. *tbx-35(-)* show 100% embryonic lethality at 25°C, significantly higher than 68% at 15°C, where the remaining 32% arrest as L1s (Fig. 2D). To determine the temporal window when temperature matters, we grew mothers at either 15°C or 25°C, shifted their embryos at the 2- or 4-cell stage to the other temperature and evaluated their development. The arrest stage tracked the temperature experienced by the mother rather than the embryo’s own post-shift temperature (Fig. 2D).

Given the maternal temperature effect, we asked how the lineage phenotypes of *tbx-35(-)* differed for embryos where mothers were grown at 25°C and 15°C. We saw earlier cell-cycle delays at higher maternal temperature. The MS4 divisions were already significantly delayed at 25°C compared with wild type but delays did not occur until MS8 at 15°C (Fig. 2F). At later stages the magnitude of delay did not differ between temperatures (15°C *vs* 25°C not significant at MS8, MS16 or MS32; Fig. 2F). We found no evidence that the earlier delay onset at 25°C reflects a difference in residual CEH-51 as the *tbx-35*-independent CEH-51::GFP remaining in mutant embryos was indistinguishable between maternal temperatures across all MS sublineages and stages (Fig. S3F). Residual CEH-51 also did not predict delay severity. Embryos with more residual CEH-51 were not those with milder cell-cycle delays (Fig. S4). Because embryos were mounted at the 2-6 cell stage and imaged at a common 22°C, the MS divisions occurred at the same temperature in both groups. The earlier onset therefore reflects a difference that is set before MS specification rather than temperature acting on the dividing cells and is independent of residual CEH-51 expression.

Together, these observations place MS specification under redundant zygotic and maternal control. The stage of embryonic arrest in tbx-35 mutants appears to be determined by the mother’s rearing temperature. Because neither the level nor the dynamics of residual CEH-51 scale with temperature, the temperature-sensitive step acts upstream of the MS lineage and in parallel to ceh-51. Its contribution surfaces only when *tbx-35* is removed, indicating that a maternally deposited, environmentally tuned factor normally buffers MS specification, and that *tbx-35* loss exposes the embryo’s latent dependence on it.

### *ceh-51* mutants have subtle, anterior-biased lineage defects

Past work reported that unlike *tbx-35* mutants, *ceh-51* mutants arrest late (often elongating or hatching) and have weaker terminal-fate defects including roughly half the number of coelomocytes, ∼25% fewer pharyngeal muscle cells, and unchanged body-wall muscle counts although some anterior muscles are mispositioned^16^. Given the asymmetrical persistence of CEH-51::GFP in anterior MS sublineages, we asked whether those lineages are preferentially disrupted in *ceh-51* mutants.

We used 4D imaging and StarryNite to trace lineages of embryos homozygous for the putative null allele *ceh-51(tm2123)*, hereafter referred to as *ceh-51(-)*. *ceh-51(-)* embryos had division-timing defects that were later, milder, and more lineage-specific than in *tbx-35* mutants (Fig. 3A, S5A). The earliest were ∼5–8 min delays in MSxpa and MSxpp at MS8; at MS16 only MSxpap was delayed (Fig. S5A). The clearest change was at MS32, where wild-type cells normally divide asynchronously across a range of times. In *ceh-51(-)* mutants they divided far more synchronously with the fastest divisions slowing and the slowest speeding up, leading to an overall reduction in the standard deviation of cell cycle length (p ≤ 0.007; Fig. 3B). Their interquartile range, similarly, narrowed from 24 to 15 min (∼40%). Finally, MSpaapp, a cell that dies in the wild type, instead divided in 14 of 19 *ceh-51(-)* embryos. Cell position defects in *ceh-51(-)* were more limited to the MS lineage and less severe overall than in *tbx-35(-)*, prominently including anterior mispositioning of MSxp-derived muscle progenitors.

**Figure 3.**
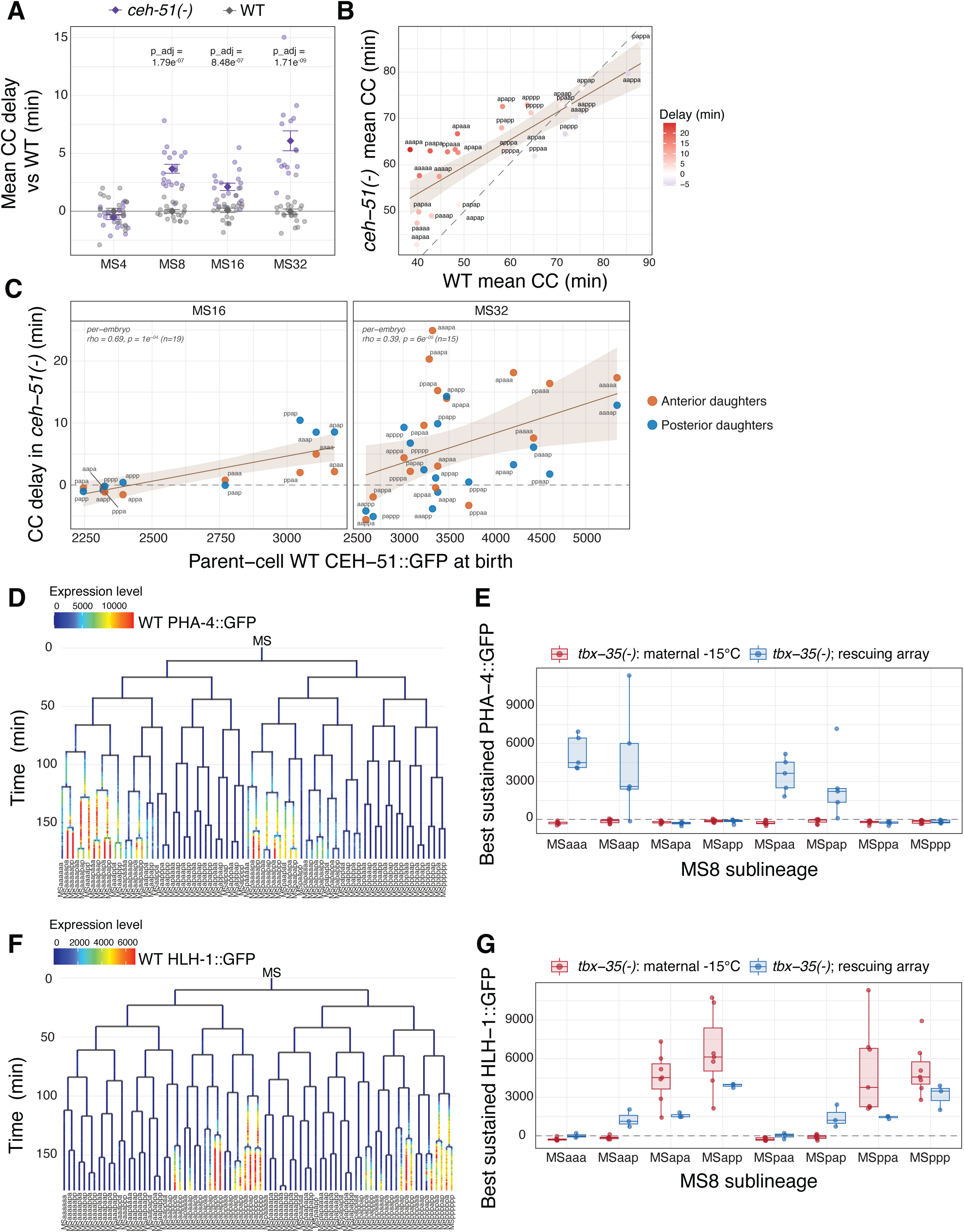
*ceh-51(-)* have later, milder, and anterior-biased defects. (A) Mean cell-cycle delay of MS divisions in *ceh-51(-)* (purple) vs wild type (grey) by stage. Diamond and bars are mean ± SE. n = 19 *ceh-51(-)*, n = 22 wild type. p values from Wilcoxon rank-sum, BH-corrected. (B) Mean cell-cycle length of each MS32 cell in *ceh-51(-)* vs wild type; each point is one cell, colored by mutant delay. Brown line, linear fit ± 95% band; dashed line, identity. n = 19 *ceh-51(-)*, n = 22 wild type. (C) Cell-cycle delay of *ceh-51(-)* MS16 and MS32 cells vs the parent cell’s wild-type CEH-51::GFP at birth. Points are individual cells, colored by anterior (orange) and posterior (blue) daughter; brown line, linear fit ± 95% band; dashed line, no delay. Per-embryo Spearman rho tested against zero by one-sample Wilcoxon: MS16 rho = 0.69 (n = 19), MS32 rho = 0.39 (n = 15). (D) Representative *tbx-35(-)*; rescuing array lineage tree (MS root) colored by PHA-4::GFP, showing the normal anterior, pharyngeal pattern. (E) Best sustained PHA-4::GFP (highest late-timepoint intensity) per MS8 sublineage in *tbx-35(−)* at 15°C (red) and *tbx-35(-);* rescuing array (blue, pooled across rearing temperatures). Boxes are median/IQR, points are per-embryo. n = 7 *tbx-35(−)*, n = 5 *tbx-35(-);* rescuing array. p values from Wilcoxon rank-sum, BH-corrected. Rescue restores PHA-4 in the pharyngeal sublineages where the mutant sits at background. (F) Representative *tbx-35(-);* rescuing array lineage tree (MS root) colored by HLH-1::GFP, showing the posterior, muscle pattern. (G) Best sustained HLH-1::GFP per MS8 sublineage, same groups. n = 7 *tbx-35(−)*, n = 3 *tbx-35(-);* rescuing array. In contrast to PHA-4, HLH-1 is higher in the mutant muscle sublineages (ectopic body-wall muscle from the MS-to-C transformation) and is reduced toward normal by the array.

Given the anterior bias of CEH-51 expression, we asked if the transcription factor preferentially regulates cell division where its levels are the highest. Since GFP matures over roughly a cell cycle at these stages, the CEH-51::GFP present in a cell reflects mostly protein inherited from its mother rather than new transcription and translation within the cell itself. We therefore related each cell’s division delay in *ceh-51(-)* mutants to its parent’s CEH-51 level. Cells whose mother had higher CEH-51::GFP were more delayed than daughters of cells with low CEH-51::GFP at both MS16 and MS32 (Fig. 3C). CEH-51::GFP in the dividing cell (as opposed to the mother) does not predict division timing defects; the time lag is consistent with CEH-51 targets influencing cell cycle length. *ceh-51*’s contribution to division timing therefore appears to be graded and correlated with CEH-51 dose - strongest in the anterior sublineages that inherit the most CEH-51, rather than an all-or-none requirement.

### Mutations in *tbx-35* and *ceh-51* alter expression of lineage-specific fate regulators

The earliest MS fate asymmetry distinguishes the pharyngeal MSxa granddaughters (which express *pha-4*) from the myogenic MSxp granddaughters (which express *hnd-1*, then *hlh-1* and *unc-120,* Fig. 1A, 3D). Prior characterization found fewer terminal PHA-4 or CEH-22-positive pharyngeal cells in *tbx-35* mutants but did not determine whether early lineage-specific expression is altered^17^. We measured lineage-resolved PHA-4::GFP (fosmid reporter) in *tbx-35(-)* embryos imaged at 22°C, after their mothers were reared at either 15°C or 25°C. Expression was greatly reduced across the pharyngeal progenitors MSaaa, MSaap, MSpaa and MSpap (Fig. 3E, S5B,F). Strikingly, residual PHA-4::GFP expression was temperature-dependent, appearing in 3 of 9 embryos at 25°C (always including the MSaaa lineage; one also in MSpaa), but none of 7 embryos whose mothers were grown at 15°C (Fig. S5B). This is the opposite of the temperature trend for terminal pharyngeal-cell loss reported previously^16^. Whether this reflects differences in experimental temperature conditions (23°C vs 25°C, stable *vs* shifted temperature), the reporter (low copy integration vs extrachromosomal array), or other factors, is unclear.

The MSxp lineage in wild-type embryos produces body-wall muscle from a subset of posterior descendants, and only those clonal muscle lineages detectably express HLH-1(MyoD)::GFP, whereas the myogenic C lineage expresses HLH-1::GFP more broadly (Fig. 3F)^22^. In *tbx-35(-)*, HLH-1::GFP was expressed throughout the MSxp lineages, varying between a pattern similar to the normal MS-like pattern (posterior-restricted) and a broader C-like pattern (Fig. 3G, S5D,F), which is consistent with a partial MS to C shift. This range was seen at both maternal temperatures, with a possible trend toward broader, more C-like expression at 25 °C (Fig. S5D, F). However, embryos overexpressing *tbx-35* from a rescuing array transiently expressed HLH-1::GFP throughout the early MS lineage (MS4/MS8/MS16) in cells that don’t have detectable HLH-1::GFP in wild-type embryos, indicating that elevated TBX-35 can drive transient HLH-1 expression normally seen only at low levels at this stage (Fig. 3G, S5F).

Consistent with the more subtle fate defects observed in *ceh-51(-)* embryos, early expression of PHA- 4::GFP and HLH-1::GFP were largely normal in the absence of *ceh-51* (Fig. S5C,E,F). However, at later stages PHA-4::GFP was inappropriately maintained in the daughters of MSpappa in *ceh-51* mutants. These cells turn off PHA-4 and produce muscle in wild-type embryos (Fig. S5C). Overall levels of HLH-1::GFP were lower in *ceh-51(-)* compared with *tbx-35(-)* and we observed minor differences in reporter patterns including occasional expansion of HLH-1::GFP into inappropriate anterior lineages (e.g. MSaaapp in one embryo, MSpaapx in two others; Fig. S5E). Together, these results suggest that *tbx-35* plays a major role in distinguishing MSxa and MSxp, while *ceh-51* has subtler roles in later fate refinement or maintenance, including occasional inappropriate reporter expression in anterior sublineages.

### *tbx-35* is required for a downstream Notch induction to the AB lineage

Given the pleiotropic phenotypes of *tbx-35* mutants, we asked whether *tbx-35* loss disrupts the ability of MS descendants to provide a Notch signal to the AB lineage. MS-derived cells drive a series of Notch inductions. The first, at the 16-cell stage, specifies the ABalp/ABara fates that make the AB pharynx^23^. This induction appeared normal in all *tbx-35(−)* embryos, where the Notch-dependent difference in cell-cycle length between each induced AB cell and its uninduced sister was indistinguishable from wild type at both maternal temperatures, and PHA-4::GFP was expressed in its normal pattern in the AB-derived pharyngeal lineages (ABalp and ABara), indistinguishable from *tbx-35(+)* rescue (Fig. 4A,B). This is consistent with past studies showing AB pharynx is present in *tbx-35(-)*, *tbx-35*;*ceh-51(-)* and *med-1;med-2(-)* mutants but not in *skn-1*(maternal), which suggests the relevant signal is a SKN-1 target acting in parallel to TBX-35 and CEH-51^16^.

**Figure 4.**
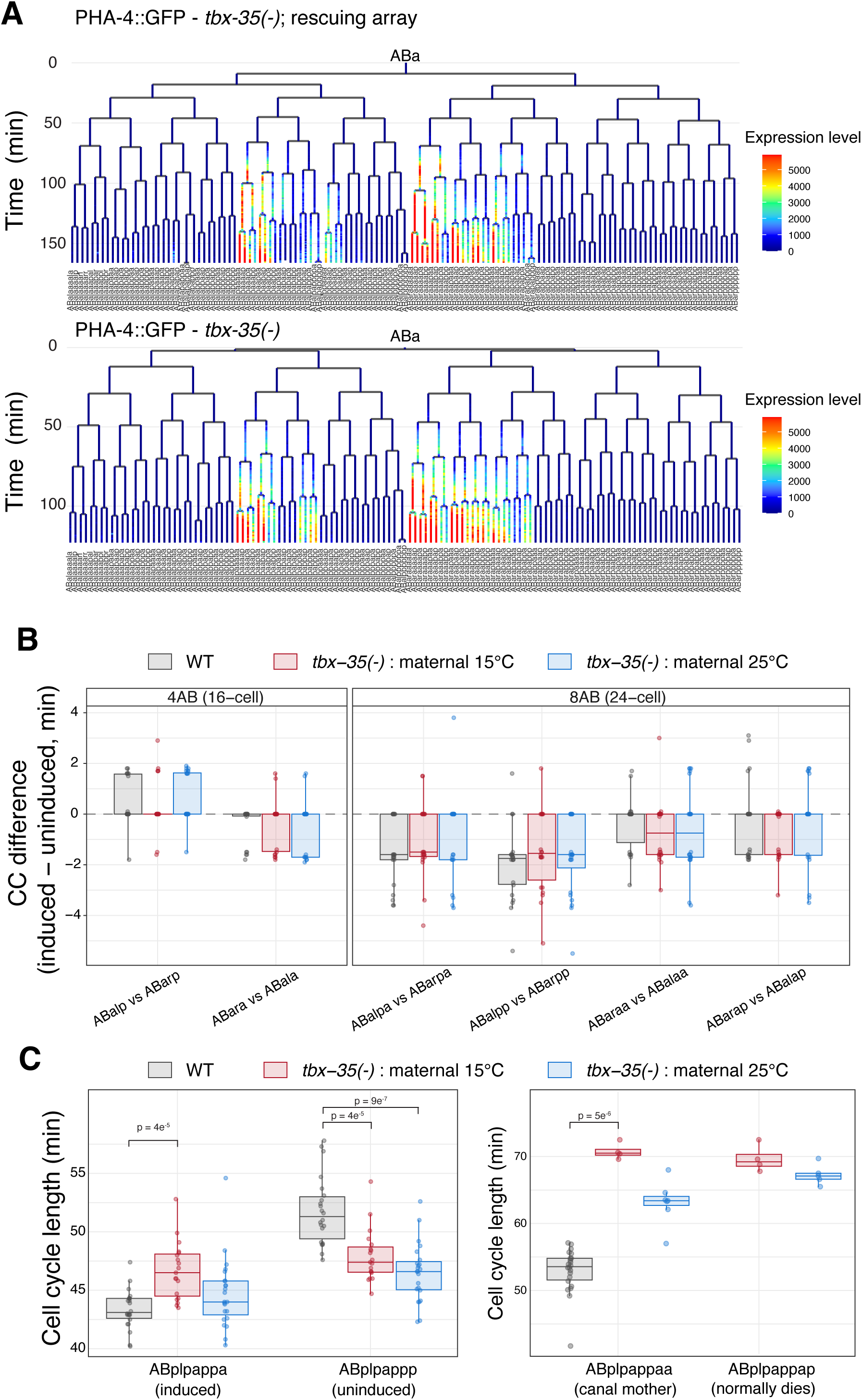
MS-derived Notch inductions in the AB lineage. (A) PHA-4::GFP in the ABa lineage (the MS→ABa Notch induction of anterior pharynx), in *tbx-35*(-); rescuing array (top) and *tbx-35*(−) (bottom); ABa root, branch color is PHA-4::GFP, cell names at the leaves. Both embryos are shown to a common 150-min window; the latest-onset pharyngeal cells (ABalpp branch) are only beginning to express at this point in both. Representative single embryos. (B) Cell-cycle asymmetry (induced − uninduced sister) for AB Notch pairs at the 16-cell (4AB) and 24-cell (8AB) stages, in wild type (grey), *tbx-35*(−) maternal 15 °C (red), and *tbx-35*(−) maternal 25°C (blue). Boxes are median/IQR. (C) Cell-cycle length of ABplpappa (induced) and ABplpappp (uninduced) [left], and of ABplpappaa (excretory-canal mother) and ABplpappap (its normally-dying sister) [right], same three groups. p values from Wilcoxon rank-sum. n = 22 wild type, n = 22 *tbx-35*(−) maternal 15 °C, n = 24 *tbx-35*(−) maternal 25 °C.

MS-derived Notch signaling to an AB-derived cell on the left side (ABplpapp) but not its corresponding cell on the right side (ABprpapp) allows the induced cell to produce several asymmetric cell fates, including the excretory canal (ABplpappaap), a cell death (ABplpappap) and certain rectal cell types^24^. We found that in *tbx-35(-)* embryos the parent and grandparent of the excretory canal cell (ABplpappa and ABplpappaa), which normally have short cell cycles, each divided ∼13 minutes later than in wild-type embryos (Fig. 4C). The sister of the excretory canal cell normally undergoes programmed cell death but instead divided in 9 of the 11 embryos tracked late enough to score. These changes made the ABplpapp lineage resemble the non-induced ABprpapp pattern and suggest that the Notch induction of ABplpapp is defective in *tbx-35* mutant embryos.

### Identification of *ceh-51* and *tbx-35* targets expressed in the early MS lineage

To determine whether *tbx-35* and *ceh-51* regulate shared or distinct targets, and whether they preferentially regulate targets at specific times or in specific sublineages of the MS lineage, we performed single-cell RNA-seq of early embryonic cells from each mutant. For each mutant, we analyzed a homozygous strain carrying a rescuing (*tbx-35+* or *ceh-51+*) transgene that is lost sporadically in progeny. We scored cells as mutant or rescued using additional markers included on the rescuing transgene (ubiquitous *eft-3promoter::mNeonGreen* and MS lineage-specific *tbx-35promoter::HIS-24::mCherry*) that could be identified in the single-cell data. We integrated the mutant and rescued cells with existing lineage-annotated wild-type reference data, clustered subsets comprising cells from early lineages, and used known markers to identify early cells from the MS and other early lineages (Fig. 5A-C).

**Figure 5.**
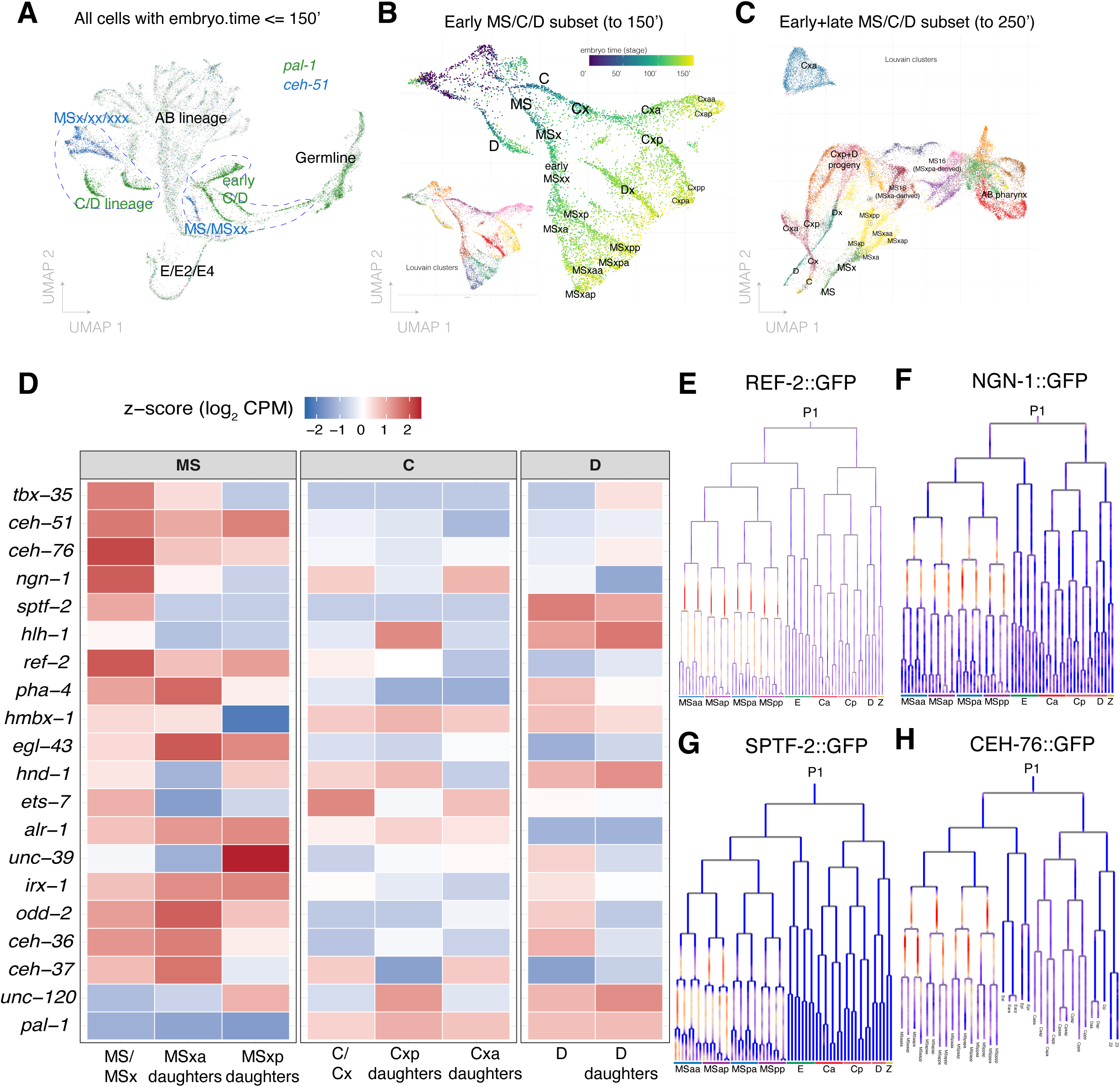
single-cell RNA-sequencing data and identification of early MS-lineage transcription factors. (A-C) UMAP projections of the CCA-integrated mutant, rescued and wild-type reference cells for (A) all cells with inferred embryo times <= 150 minutes (∼100-cell stage) (B) a subset of the cells in A from the early MS, C and D lineages and (C) MS, C and D-lineage derived cells from embryos inferred to be <= 250 minutes (∼350-cell stage). (D) enrichment of selected TFs in clusters corresponding to the listed lineages (MS/MSx, MSxa, etc) compared to other early cells at that stage, showing lineage-specificity of each factor. (E–H) Reporter protein expression: (E) REF-2::GFP (wgIs520), (F) NGN-1::GFP (nIs394), (G) SPTF-2::GFP (wgIs769), (H) CEH-76::GFP (ceh-76(ot1042)).

Because earlier studies and this work suggest a partial MS-to-C transformation in *tbx-35* mutants, we first asked whether early MS lineage cells could still be recognized from their transcriptomes in each mutant. We identified clusters corresponding to MS or the MS daughters (MSx), granddaughters (MSxx) or great-granddaughters (MSxxx) (Fig. 5B,C), and compared the frequency of mutant vs rescued cells in these clusters. We found no significant difference in cell frequencies within the MS lineage clusters overall, or in clusters corresponding to the MSx (MS daughters), MSxx and MSxxx stages, suggesting that most of the mutant MS lineage cells can be annotated (Fig. 5A, S6A). Most *tbx-35* mutant cells continued to express *ceh-51* mRNA, consistent with our lineage tracing results (Fig. 6B). Other EMS lineage specific factors such as *sdz-1* also retained expression, but many other early MS lineage markers were reduced.

**Figure 6.**
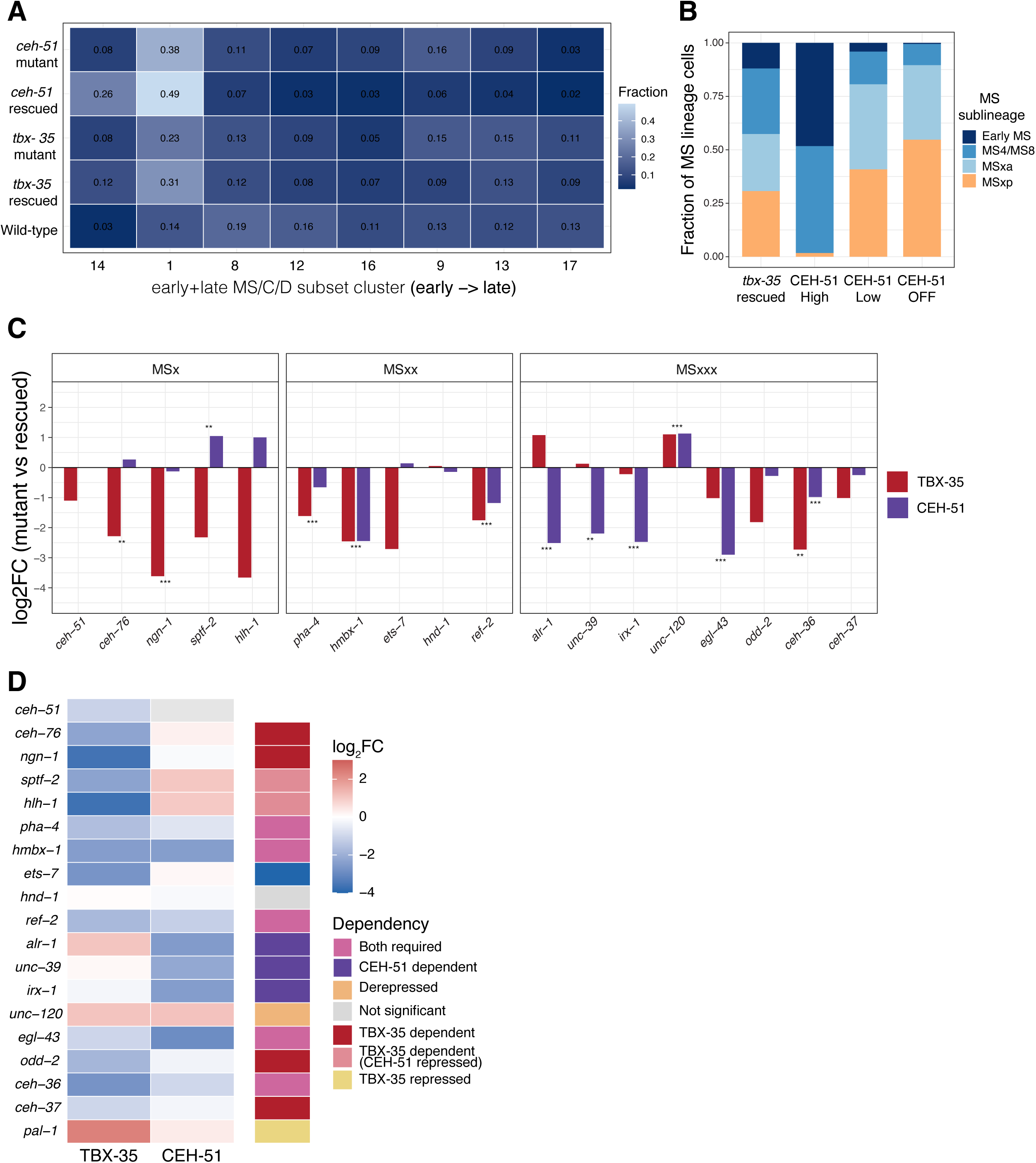
Identification of expression defects in *tbx-35* and *ceh-51* mutant cells. (A) Fraction of cells from each annotation (rescued, mutant, wild-type) in each cluster (250’ MS/C/D subset, see Fig. 5C). (B) fraction of *tbx-35* mutant or rescued cells in each MS lineage subcluster, subdivided by *ceh-51* expression status. (C) Fold-change in three divisions of the MS lineage for identified targets, comparing the effect size of *tbx-35* vs *ceh-51* mutants. In these comparisons, the DEG calculations were done only on the noted lineage subset (e.g. MSx or MSxx) even if the target gene normally is expressed across multiple stages (D) Categorization of *tbx-35* and *ceh-51* targets.

To interpret mutant expression defects, we first defined the normal regulatory landscape of the early MS lineage by identifying transcription factors enriched in MS-derived cells relative to contemporaneous cells from the AB, C, D, and E lineages. This recovered known stage- and sublineage-specific regulators, including *med-1/2, tbx-35*, *ceh-51*, *pha-4,* and *hnd-1*, along with additional MS-enriched factors with less-characterized embryonic roles (19 TFs in total; Figure 5D). Several of these were validated by 4D imaging and StarryNite lineage tracing (Fig. 5E-H).

Overall, our analysis identified distinct, partially overlapping, targets for *tbx-35* and *ceh-51* (Fig. 6C,D, S6B,C). Targets of *tbx-35* were biased towards the earlier stages of MS development; of the five TFs initially expressed in the MS daughters, three (the novel homeodomain protein *ceh-76, ngn-1/Neurogenin* and *sptf-2/Sp2*) had strong and significant decreases in *tbx-35* mutants (log2fc > 2; q <= 0.05). *hlh-1/MyoD* had very weak expression at this stage in wild-type embryos and was also possibly reduced (albeit not significantly) in *tbx-35(-)*, while *ceh-51* had only a modest, non-significant decrease (20% FDR, log2fc ≈ 1). This *tbx-35*-independent residual expression of *ceh-51* is consistent with our StarryNite analysis of CEH-51::GFP and may explain the stronger phenotype of *ceh-51(-); tbx-35(-)* double mutants compared to *tbx-35(-)*. None of the MSx factors had substantial expression decreases in *ceh-51* mutant cells. *tbx-35* was also required for full expression of several MS-granddaughter expressed genes including MSxa-expressed *pha-4/FoxA* and *hmbx-1/HMBOX*, and MSxx-expressed *ref-2/Zic*.

At later stages, the dependencies diverged. *tbx-35* remained important for several MS-granddaughter regulators, whereas *ceh-51*-dependent targets became more prominent in the next generation and were biased toward lineally anterior MS sublineages. The most strongly *ceh-51*-dependent gene in the MS granddaughters was *hmbx-1*, with weaker quantitative reductions in *pha-4* and *ref-2* expression. This is consistent with our observation of substantial loss of PHA-4::GFP in *tbx-35(-)* but minimal early changes in *ceh-51(-)*. Stronger *ceh-51* defects and distinct *tbx-35*-dependent changes were seen in the next generation (MSxxx; Fig. 6C,D S6B,C). Notably the *ceh-51*-regulated targets were initially activated in lineally anterior daughter cells: MSxa (*hmbx-1*), MSxaa (*egl-43*), MSxpa (*unc-39*), MSaaa (*alr-1*) or MSpaa (*irx-1*). In contrast the *tbx-35*-dependent genes *odd-2*, *ceh-36* and *ceh-37*, were primarily expressed in the posterior lineage MSxap. These patterns suggest that the two factors have both lineage- and stage-specific roles in MS patterning and are intriguing given the anterior bias we observed for CEH-51::GFP expression. (Fig. 2).

We also tested for repressed targets (with increased expression in either mutant compared to rescued cells). In *tbx-35* mutant cells, *pal-1* was significantly upregulated throughout the early MS lineage (Fig. S6C), consistent with a partial MS-to-C lineage transformation. The muscle-promoting factor *unc-120/Srf* had higher levels in both mutants, also consistent with the earlier muscle specification (*hlh-1* expression) seen in some of the *tbx-35* mutant MSxp lineages by imaging (Fig. S5D,F). Levels of some MS-enriched genes, including the Notch ligand *lag-2*, were not significantly altered in either mutant. This is consistent with the preserved early Notch-mediated induction of AB-derived pharyngeal fates but leaves open why later MS-dependent Notch inductions are defective in *tbx-35* mutants.

Because several MSx-expressed TFs were strongly *tbx-35*-dependent, we asked whether any might act redundantly with *ceh-51*. We focused on *ceh-76*, a strongly MSx-specific homeodomain gene distantly related to *ceh-51*. However, *ceh-76(tm10678)* single mutants had no major viability defects or changes in division-timing or position defects as measured by StarryNite and *ceh-51;ceh-76* double mutants resembled *ceh-51* single mutants. Similarly, *ngn-1(ok2200)* embryos and previously characterized *ref-2(gk178)* embryos^21^ showed only limited MS-lineage division timing defects. Thus, no single tested MSx factor accounts for the strong *tbx-35* phenotype, suggesting that TBX-35 controls a distributed early regulatory program.

## Discussion

### TBX-35 coordinates multiple dimensions of MS lineage identity

Our results show that the partially redundant MS lineage regulators TBX-35 and CEH-51 make distinct contributions across developmental time, sublineage, and phenotype. Previous studies established that loss of *tbx-35* partially transforms MS toward a C-like fate^16,17^. We find that this transformation extends beyond terminal cell identity to the dynamic behavior of the lineage itself. Cell-cycle timing is itself highly reproducible and linked to lineage and fate in the *C. elegans* embryo, and previous fate transformations can be accompanied by corresponding changes in division timing and pattern^20^. *tbx-35* mutant MS cells divide more slowly, increasingly approach the characteristic timing of homologous C-lineage cells, and acquire aspects of the C-specific pattern. At the same time, the transformation is incomplete - mutant MS cells do not simply adopt the positions of the corresponding C descendants, and several fate and signaling phenotypes remain intermediate. These observations argue that lineage identity is not a single binary state but a collection of partly separable properties, including cell-cycle timing, spatial behavior, transcriptional state, signaling competence, and terminal fate. Other lineage-specific transcription factors in the embryo similarly regulate progenitor division and migration phenotypes that are not apparent from terminal cell-type analysis alone^25^.

This modular view of lineage identity may explain why developmental transformations are often partial. TBX-35 loss strongly affects some components of the MS program while leaving others intact or only partially altered. For example, MS-derived cells acquire C-like division timing and broaden HLH-1 expression, yet their spatial trajectories do not simply reproduce those of C cells. The expansion of HLH-1 in *tbx-35* mutants is consistent with derepression of the C-lineage regulator PAL-1, which directly activates *hlh-1* in posterior embryonic muscle lineages^22^. Similarly, the earliest MS-dependent Notch induction into the AB lineage remains intact, whereas a later MS-dependent induction is defective^23,24^. It appears that different outputs of MS specification can persist for different lengths of developmental time after the initiating regulatory program is perturbed. Developmental transformation is therefore better viewed as a multidimensional shift in lineage state than as an all-or-none conversion from one lineage identity to another.

Finally, the nonautonomous defects in *tbx-35* mutants emphasize that lineage specification affects the developmental environment experienced by other cells. The earliest MS-dependent Notch induction of AB pharyngeal fate is preserved, whereas a later induction into the ABplpapp lineage is frequently lost. This suggests that signaling competence is acquired progressively by MS descendants and can therefore be selectively disrupted downstream of an early specification defect. More generally, mutations in lineage-specifying transcription factors may propagate through an embryo both cell-autonomously, by changing the behavior and fate of their descendants, and nonautonomously, by altering the signals those descendants provide to neighboring lineages.

### TBX-35 and CEH-51 coordinate distinct temporal and sublineage-specific programs

TBX-35 and CEH-51 also illustrate how partially redundant transcription factors can overlap without acting as interchangeable backups, as observed for other redundant transcription-factor pairs in *C. elegans* development^26,27^. TBX-35 has the broader and earlier role where its loss disrupts early MS regulators, division behavior, lineage-specific fate markers, and later intercellular signaling. CEH-51 loss produces later, milder, and more sublineage-restricted phenotypes. CEH-51 protein itself becomes enriched in anterior daughters over successive MS divisions, and cells descended from mothers with the highest normal CEH-51 levels show the strongest division-timing defects when *ceh-51* is removed. The single-cell data parallel these phenotypic differences, and we find that TBX-35-dependent genes are enriched at earlier stages, whereas CEH-51-dependent genes become more prominent later and are biased toward lineally anterior MS sublineages. Together, our results suggest that redundancy between TBX-35 and CEH-51 is temporally and spatially structured rather than uniform.

Our scRNA-seq data further suggest that TBX-35 controls a distributed regulatory program. Several transcription factors expressed shortly after MS specification depend strongly on TBX-35, yet loss of individual tested factors such as *ceh-76* or *ngn-1* produces only limited lineage phenotypes. This argues against the broad *tbx-35* phenotype being mediated through a single downstream regulator. Instead, TBX-35 may establish an early regulatory state through multiple partially overlapping effectors whose combined activities control fate specification, cell-cycle timing, positioning, and signaling. A distributed regulatory architecture has also been described downstream of PAL-1 in the C lineage, where downstream regulators are activated in successive temporal phases and contribute to specification, differentiation, and morphogenesis^19^. Distinguishing direct from indirect regulation will require cis-regulatory and binding analyses, but the lineage-resolved mutant transcriptomes provide a framework for identifying the combinations of downstream regulators that generate particular developmental phenotypes. The rescuing-array strategy used here also provides an internally controlled way to profile embryonic lethal mutants by single-cell RNA-seq, because mutant and rescued cells are collected and processed within the same samples.

### Maternal inputs provide an additional layer of robustness to MS specification

The temperature dependence of *tbx-35* mutants reveals a further source of developmental robustness. Embryonic outcome depends strongly on the temperature experienced by the mother and shifting embryos after the 2–4-cell stage does not reset this effect. The earlier onset of MS division defects in embryos from mothers grown at high temperature is therefore determined before the MS lineage is specified. This difference is not explained by residual CEH-51 abundance or dynamics, and residual CEH-51 does not predict the severity of the cell-cycle phenotype. These observations are consistent with a maternal or very early embryonic activity that buffers MS development and becomes phenotypically important when TBX-35 is absent. Such cryptic buffering mechanisms may be difficult to detect in wild-type embryos because their effects emerge only after a major regulatory input is compromised.

In conclusion, our results support a model in which developmental robustness emerges from overlapping but non-equivalent regulatory inputs. TBX-35 establishes a broad early MS program, CEH-51 reinforces and refines selected later and anterior components of that program, and additional maternal activity provides a further layer of buffering. This organization allows the MS lineage to maintain robust development while still permitting individual regulators to control distinct temporal, spatial, and phenotypic outputs.

## Methods

### *C. elegans* growth and maintenance

*C. elegans* strains were maintained at standard growth temperatures (15°C, 20°C, 22°C, or 25°C as indicated) on OP50 *E. coli* on NGM plates. *tbx-35* and *ceh-51* rescuing array strains were generated by injecting using a mix of *ceh-51*(+) in plasmid pCFJ1202, *ptbx-35*::*H1-mcherry*, *peft-3*::mNeonGreen(NLS)::*tbb-2* 3’UTR for *ceh-51* and *tbx-35*+, *tbx-35*::*H1-mcherry*, *peft-3*::mNeonGreen(NLS)::*tbb-2* 3’UTR for *tbx-35* into N2 with coinjection marker myo-3::mCherry (pCFJ104) and pBluescript. Array-positive worms were selected and after transmission was confirmed, males were generated and crossed into *tbx-35 (tm1789)* and *ceh-51(tm2123)* deletion strains respectively. Homozygous strains were confirmed by PCR with primers that distinguished array versus endogenous constructs.

### Viability experiments

Embryos were dissected from hermaphrodites of the appropriate genotypes grown under the indicated conditions, and 2–6-cell-stage embryos were transferred onto a glass slide with egg buffer containing methyl cellulose with 20 μm beads and the coverslip sealed with Vaseline. Slides were incubated at the appropriate temperature and stage of arrest and array status were scored the next day. For temperature shifts, L4s were moved to the appropriate temperatures and embryos incubated at the alternate temperature as specified for the experiment.

### 4D imaging, lineage tracing and reporter expression imaging

We obtained confocal time-lapse images using a Leica Stellaris laser scanning confocal microscope (67 z-planes at 0.5 μm spacing and 1.5 min time spacing, with laser power increasing by 4-fold through the embryo depth to account for attenuation of signal with depth). Embryos were dissected from hermaphrodites of appropriate genotypes and mounted in egg buffer/methyl cellulose with 20 μm beads used as spacers and imaged at 22 °C using a stage temperature controller^9^. We used StarryNite software to automatically annotate nuclei and trace lineages^11^. We corrected errors from the automated analysis and quantified reporter expression in each nucleus relative to the local background (using the “blot” background correction technique) with AceTree software as previously described^6,12^.

### Cell lineage analysis and cell-cycle timing

Lineaged embryos were corrected in AceTree and exported as per-nucleus tables (CD/ACD files) containing, for each tracked nucleus, its lineage name, time, xyz position, and background-corrected reporter intensity. All downstream analysis was performed in R [version 4.5.2] with custom scripts as described previously^21,25,28^. For each cell we defined its cell-cycle length as the interval between its birth and its division, taken on the normalized developmental timescale of the reference tables, which places embryos imaged under different conditions on a common developmental clock. A wild-type reference was assembled from the tracked wild-type embryos from Richards et al.^29^. For every cell present in at least five wild-type embryos we computed the mean wild-type cell-cycle length. Cell-cycle delay for a mutant cell was defined as its cell-cycle length minus the wild-type mean for the same wild-type cell (by lineage ID), in minutes. A per-embryo delay at a given stage (MS4, MS8, MS16, MS32) is the mean delay across that embryo’s cells at that stage.

### MS-to-C timing comparison

To test whether *tbx-35(-)* MS cells adopt a C-like division program, each MS cell was paired with its lineage-homologous C cell (matched by suffix within the sublineage). We used two complementary measures. (i) Gap closure: each MS cell’s delay was expressed as a fraction of the wild-type MS-to-C cell-cycle-timing gap, where 0% is wild-type MS timing and 100% is homologous wild-type C timing; per-stage values are means across cells, with 95% confidence intervals from 2000 bootstrap resamples of embryos (wild type and mutant resampled independently). (ii) Per-embryo rank correlation: for each embryo we computed the Spearman correlation between its MS cell-cycle lengths and the homologous wild-type C cell-cycle lengths, requiring at least six paired cells per embryo.

### Reporter expression quantification and depth correction

Reporter intensity in each nucleus was measured relative to local background using the “blot” method in AceTree^12^. To correct for depth-dependent signal attenuation, we fit an exponential model of expression versus imaging depth (z) to wild-type MS-lineage CEH-51::GFP over the first 150 min after MS birth and divided each measurement by the fitted depth factor and by a depth-dependent excitation term, rescaled to the mean; the same correction was applied to mutant and rescue embryos. Anterior/posterior asymmetry was computed per division as log2(anterior daughter / posterior daughter) intensity at birth. Expression at birth was taken as the mean of the first three timepoints after a cell’s birth. “Best sustained” expression per MS8 sublineage was defined as the maximum, across that sublineage’s descendant cells, of each cell’s late expression (the mean over its last five tracked timepoints); taking a maximum reports whether the reporter is maintained in at least one branch of the sublineage without diluting fate-restricted expression. Residual CEH-51::GFP in *tbx-35(-)* was quantified per MS cell as the mean of its three brightest depth-corrected timepoints, relative to a per-embryo detection floor defined as the mean + 2 SD of that embryo’s own non-MS cells (AB, C, D and E), which do not express CEH-51.

### Position and nearest-neighbor analysis

Nuclear positions were compared to the mean wild-type reference positions. For each cell, at four timepoints spanning its lifetime (10, 35, 70 and 90% of its tracked duration), we identified all co-existing cells and computed the Euclidean distance from that cell to each of them, both in the mutant embryo and in the wild-type reference. At each timepoint we took the mean of the ten largest absolute log2 ratios of mutant-to-wild-type distance (the ten most-distorted of the cell’s spatial relationships); the cell’s neighbor-deviation score was the median of these four timepoint values (cells tracked for fewer than two frames were omitted). Higher scores therefore indicate a greater departure of a cell’s spatial relationships from the wild-type configuration, and a per-embryo score at each stage is the mean across that embryo’s cells. Three-dimensional renderings show aligned nuclear positions for individual representative embryos, colored by MS4 sublineage.

### Statistical analysis

Analyses were performed in R [version 4.5.2]. Unless stated otherwise, group comparisons used the two-sided Wilcoxon rank-sum (Mann-Whitney) test on per-embryo summary values, with Benjamini-Hochberg (BH) correction across the family of tests within each panel; p_adj denotes BH-corrected p values. To avoid pseudoreplication, cells were aggregated to a single value per embryo before testing, and n throughout refers to the number of embryos (given in each figure legend). Deviations from zero (cell-cycle delay, anterior/posterior asymmetry, and per-embryo correlations) were assessed with the one-sample Wilcoxon signed-rank test. Correlations are Spearman’s rho; where a correlation was summarized across embryos (e.g., residual CEH-51 vs cell-cycle delay, or MS vs C timing), the per-embryo rho values were tested against zero by one-sample Wilcoxon. Bootstrap confidence intervals used 2000 resamples of embryos. Box plots show the median and interquartile range; points are per-embryo values; error bars are mean ± SE or median ± IQR as indicated in each legend.

### Single-Cell RNA sequencing

We collected embryos by hypochlorite treatment, and dissociation into single cells using chitinase and aspiration as described previously^3^. Cells collected were mixed-stage, but we biased the samples for early-stage embryos by washing mothers off the plate for hypochlorite treatment once the first embryos were visible. scRNA-seq was performed using the 10x Genomics 3’ Gene Expression v3.1 assay, and libraries were sequenced on an Illumina NovaSeq X sequencer. We did primary cell calling using Cell Ranger software; we collected three biological replicates each per strain (PSV014 [*tbx-35(tm1789)* II; (*tbx-35+* rescuing array)] and PSV045 [*ceh-51(tm2123)* V; (*ceh-51+* rescuing array)]), totaling 72,232 newly collected cells after filtering. Data were corrected for ambient background using SoupX^30^, assigned cell-specific embryo-time (stage) estimates, and integrated with annotated wild-type reference datasets using the CCA method and Louvain clustered in the Seurat package^3,31–33^, and visualized in Viscello to annotate lineages using known markers and previously annotated cells from the reference datasets. Clusters containing early cells of the appropriate MS, C, D, E and AB lineages, further filtered by embryo.time estimates, were re-integrated and re-clustered to identify clusters corresponding to specific early lineages for analysis. Differentially expressed genes were identified in relevant clusters using FindMarkers in Seurat, by comparing cells annotated as “rescued” or “mutant.” Rescue/mutant status was based on detection of reads only present on the rescuing array, e.g. *ceh-51* or *tbx-35* (depending on the strain), mCherry or mNeonGreen. All analysis code will be made available on GitHub and raw data will be made available at the Gene Expression Omnibus (GEO).

