## Supplementary Figures for "Distinct roles for partially redundant transcription factors in *Caenorhabditis elegans* mesoderm lineage development"

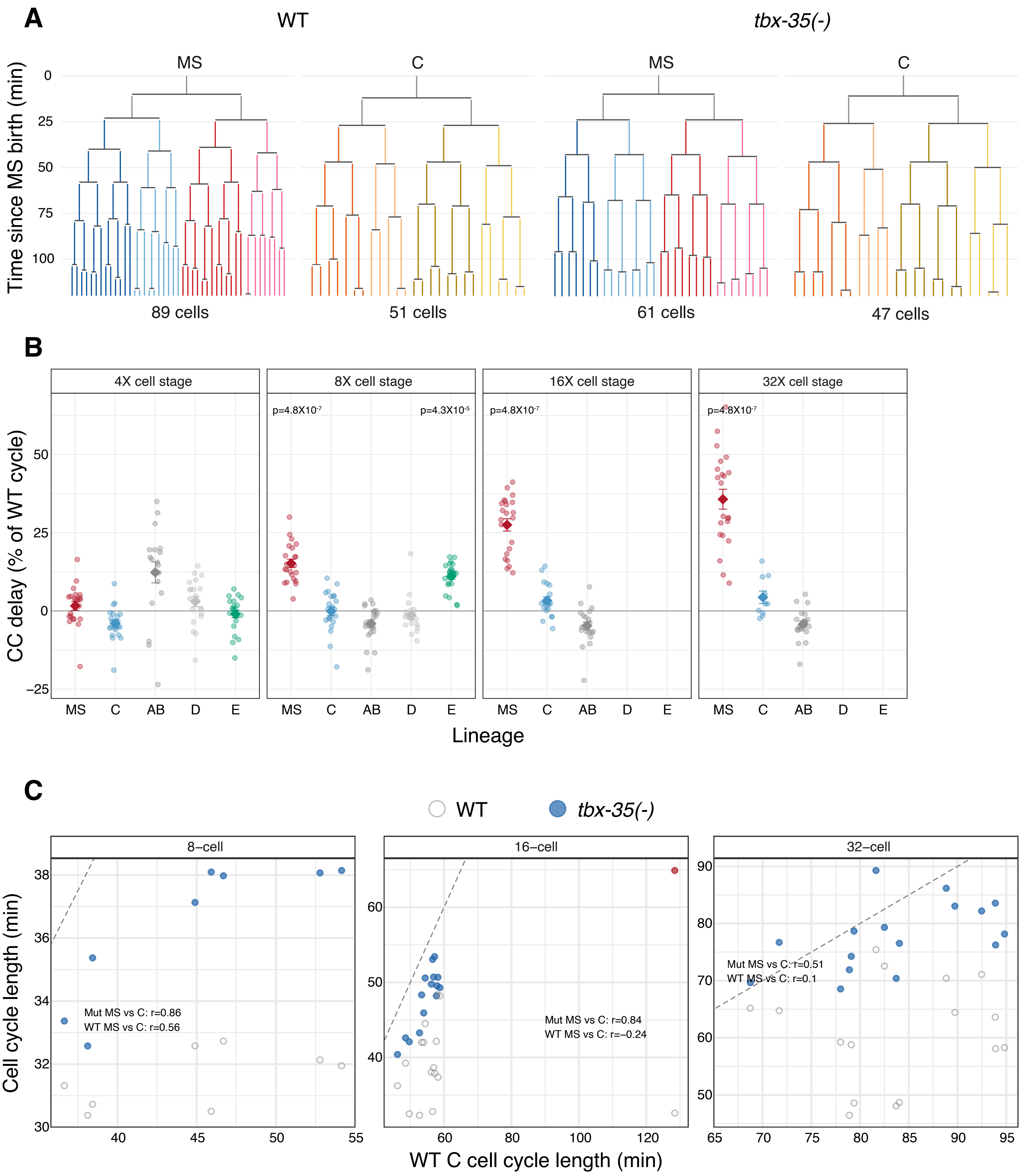


Supplementary Figure 1

**Supplementary Figure 1. Division-timing defects are specific to MS and E.**

(A) MS and C division-timing trees for a representative wild-type and a representative *tbx-35(-)* embryo, each aligned to its own MS birth (t = 0), colored by sublineage; cell counts beneath (wild type MS 89, C 51; *tbx-35(-)* MS 61, C 47). (B) Cell-cycle delay as a percentage of the wild-type cycle by lineage (MS, C, AB, D, E) at each lineage's 4-, 8-, 16- and 32-cell stage. Diamond and bars are mean ± SE (points with $n < 10$ embryos, and delays < 8 min within single-frame tracking resolution, excluded). p values from one-sample Wilcoxon signed-rank vs 0 (MS at all stages; E at the 8-cell stage). (C) Per-cell MS-C timing: each point is one MS cell, its cycle length vs the lineage-homologous wild-type C cell at the 8-, 16- and 32-cell stages; open, wild-type MS; filled, *tbx-35(-)* MS. Dashed line, identity. r values are Pearson pooled across cells (descriptive; the per-embryo test is Fig. 1D).

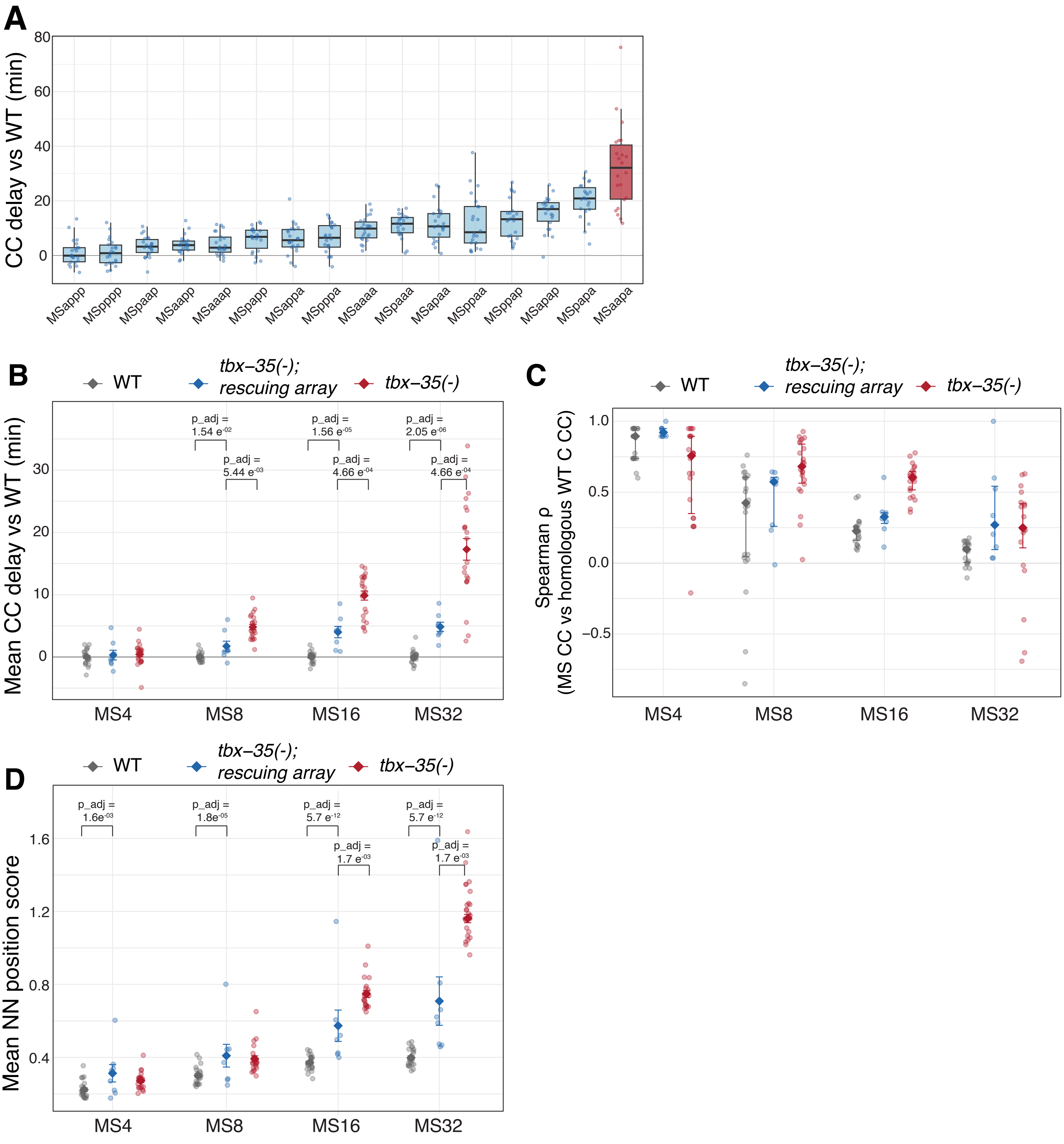


Supplementary Figure 2

**Supplementary Figure 2. Sublineage defects and rescue in *tbx-35(-)*.**

(A) Cell-cycle delay of each MS16 cell in *tbx-35(-)*, ordered by magnitude; MSaapa (red) is the most delayed, mirroring its C homolog. Boxes are median/IQR, whiskers show range. n = 22 *tbx-35(-)*. Descriptive ranking. (B) Mean cell-cycle delay for wild type (grey), *tbx-35(-);* rescuing array (blue), and *tbx-35(-)* (red) by stage. Diamond and bars are mean ± SE. n = 22 wild type, n = 8 *tbx-35(-);* rescuing array, n = 22 *tbx-35(-)*. p values from Wilcoxon rank-sum, BH-corrected. (C) Per-embryo Spearman rho of MS with homologous wild-type C cell-cycle lengths, same three groups. Diamond and bars are median ± IQR. n = 6 wild type, n = 8 *tbx-35(-);* rescuing array, n = 22 *tbx-35(-)*. The elevated C-correlation of *tbx-35(-)* is reduced toward wild type in *tbx-35(-);* rescuing array, clearest at MS16. (D) Mean nearest-neighbor position score for the same three groups by stage. Diamond and bars are mean ± SE. n = 22 wild type, n = 8 *tbx-35*(-); rescuing array, n = 22 *tbx-35(-)*. p values from Wilcoxon rank-sum, BH-corrected.

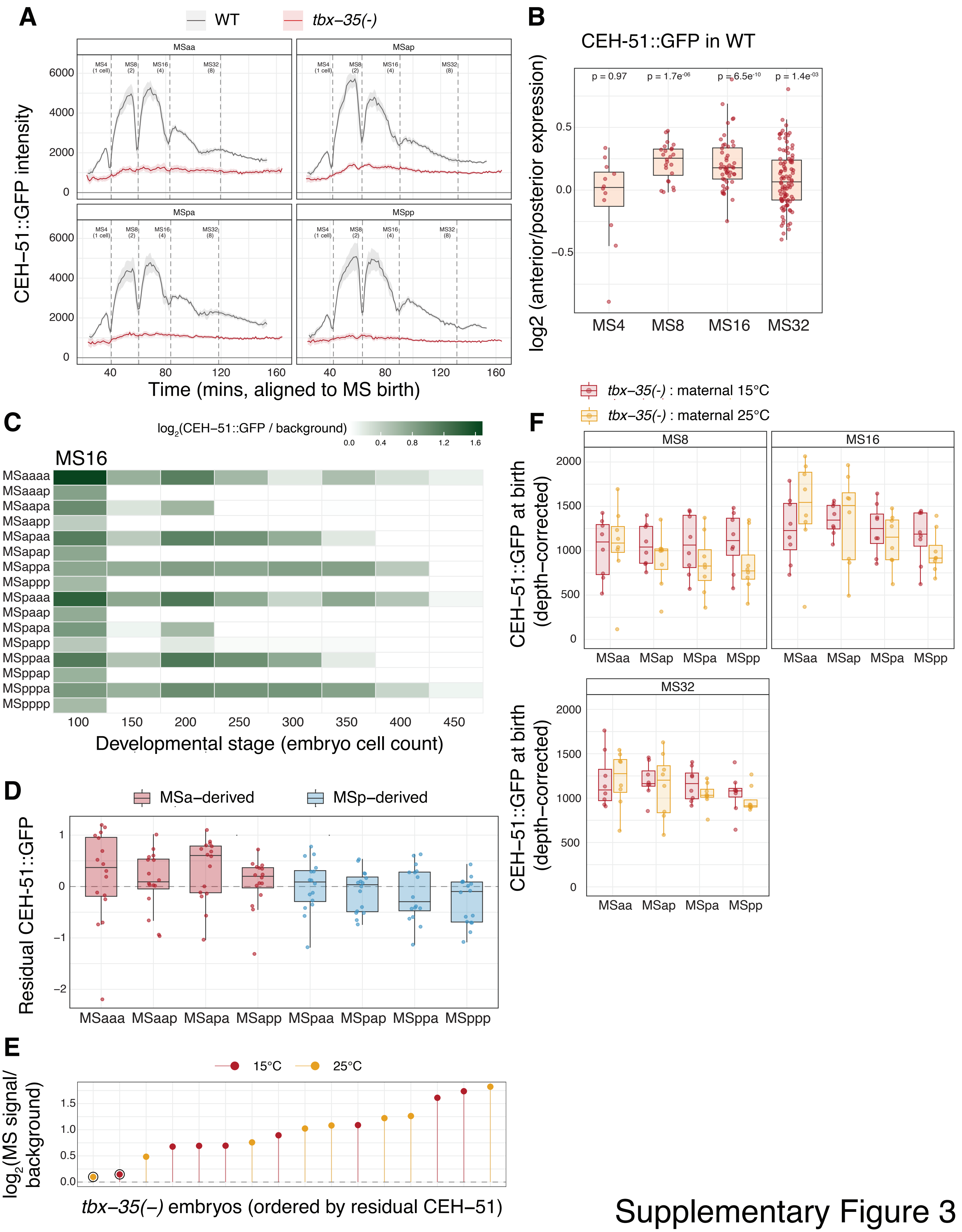


Supplementary Figure 3

**Supplementary Figure 3. CEH-51::GFP dynamics in wild type and its loss in *tbx-35(-)*.**

(A) Depth-corrected CEH-51::GFP over time (aligned to MS birth) for the four MS4 sublineages, wild type (grey) vs *tbx-35(-)* (red); line and band, mean ± SE; division times and stages marked. n = 6 wild type, n = 8 *tbx-35(-)*. (B) Per-division log2(anterior/posterior CEH-51::GFP) in wild type by stage. Boxes are median/IQR, points are divisions. n = 6 wild type. One-sample Wilcoxon signed-rank vs 0. (C) Persistence of CEH-51::GFP: log2(CEH-51::GFP / background) per MS16 sublineage (rows, anterior to posterior) by developmental stage (embryo cell count). n = 6 wild type. (D) Residual CEH-51::GFP per MS8 cell in *tbx-35(-)*, MSa-derived (red) vs MSp-derived (blue), anterior to posterior. Boxes are median/IQR. n = 16 *tbx-35(-)* (pooled across temperature). (E) Per-embryo residual CEH-51::GFP: log2(brightest MS cell / background) for each *tbx-35(-)* embryo, ordered by residual level; 15°C (red), 25°C (amber); the two ringed points were scored undetectable. n = 16 *tbx-35(-)*. (F) Mean CEH-51::GFP at birth (depth-corrected) per MS4 sublineage at MS8/MS16/MS32 in *tbx-35(-)*, from mothers reared at 15°C (red) vs 25°C (amber). Boxes are median/IQR, points are per-embryo. n = 8 per temperature. Levels did not differ between maternal temperatures at any sublineage or stage (Wilcoxon rank-sum, BH-corrected).

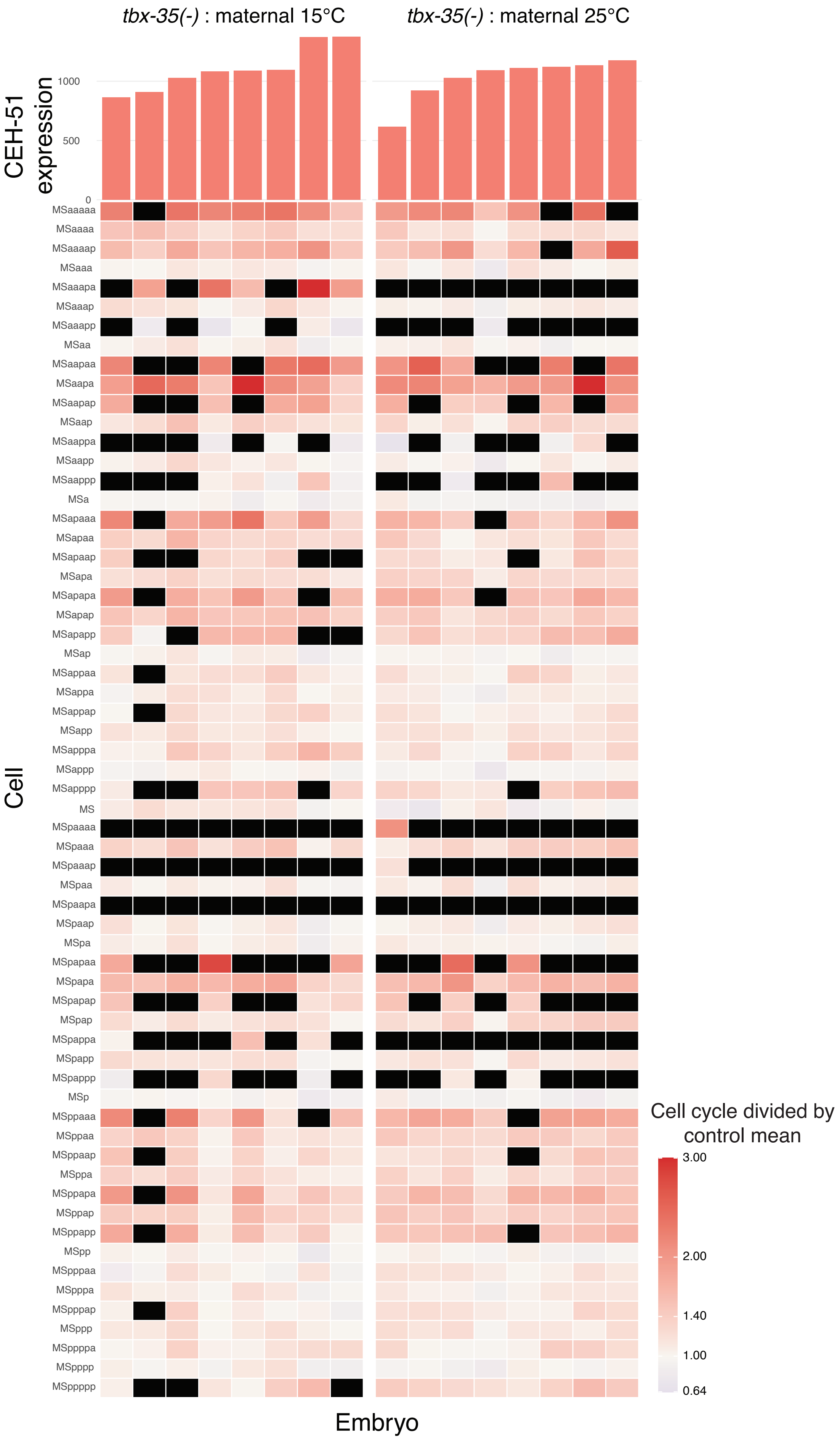


Supplementary Figure 4

**Supplementary Figure 4. Residual CEH-51 does not track cell-cycle delay in *tbx-35(-)*.**

Per-embryo CEH-51::GFP (top bar, mean MS expression) aligned above per-cell cell-cycle length (heatmap, cycle divided by the wild-type mean; white is wild-type timing, red is delayed, black is not tracked or expressed) for *tbx-35(-)* embryos from mothers at 15°C (left) and 25°C (right). Columns are individual embryos ordered by increasing expression, n = 8 *tbx-35(-)* per temperature; rows are MS cells. Per-embryo residual CEH-51 was uncorrelated with per-embryo mean cell-cycle delay (Spearman rho = 0.08, p = 0.77, n = 16; computed per embryo to avoid pseudoreplication).

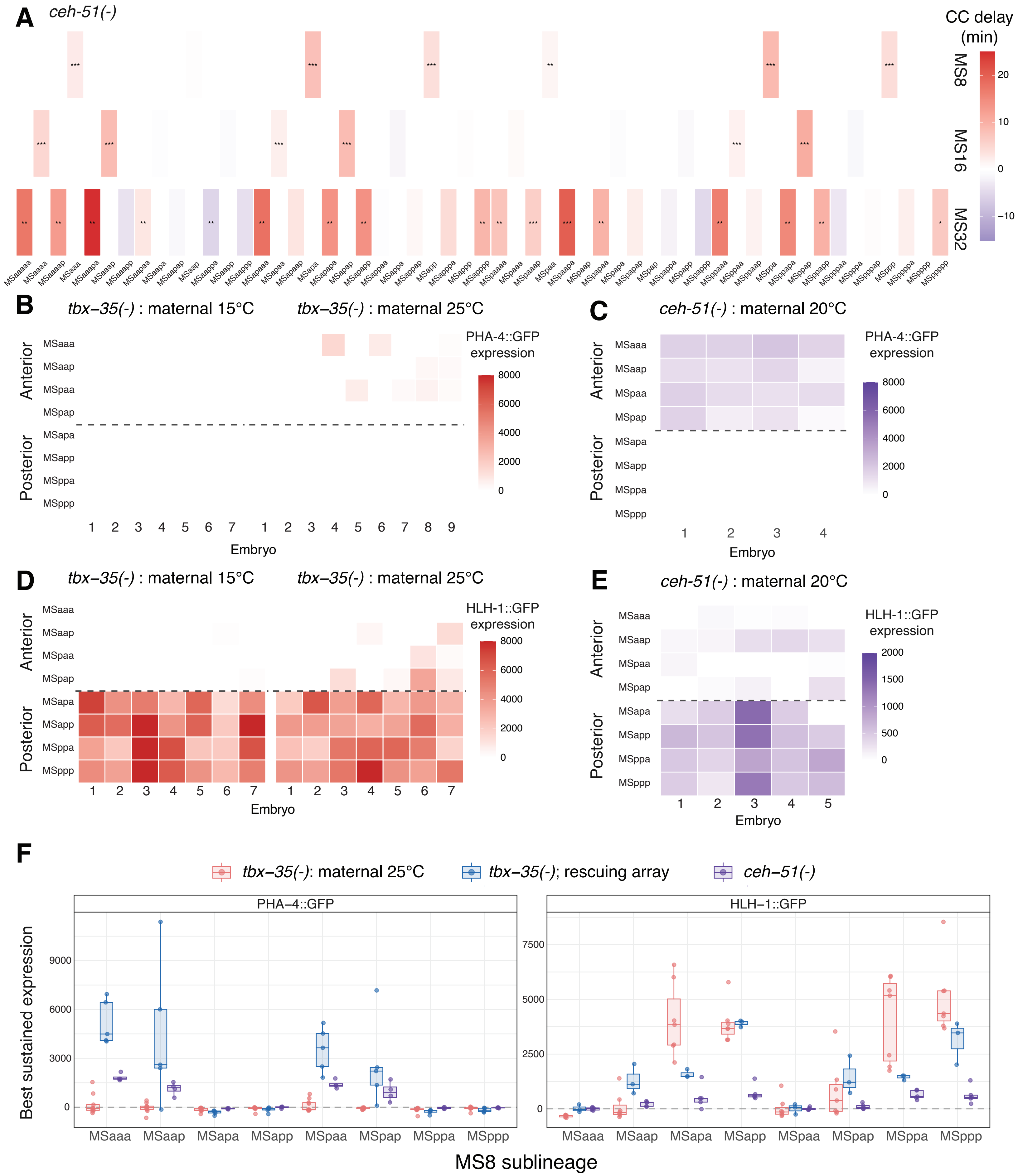


Supplementary Figure 5

**Supplementary Figure 5. Cell-cycle delay in *ceh-51(-)* and reporter expression across genotypes.**

(A) Per-cell cell-cycle delay in *ceh-51(-)* across the MS lineage at MS8, MS16 and MS32 (heatmap, minutes relative to wild type; anterior cells at left, posterior at right). Asterisks mark cells significantly delayed relative to wild type (Wilcoxon rank-sum, BH-corrected). n = 19 *ceh-51(-)*. (B) PHA-4::GFP per MS8 sublineage (anterior above, posterior below the dashed line) in each *tbx-35(-)* embryo, at 15°C and 25°C. n = 7 *tbx-35(-)* 15°C, n = 9 *tbx-35(-)* 25°C. (C) PHA-4::GFP per MS8 sublineage in each *ceh-51(-)* embryo at 20°C. n = 4 *ceh-51(-)*. (D) HLH-1::GFP per MS8 sublineage in each *tbx-35(-)* embryo, at 15°C and 25°C. n = 7 *tbx-35(-)* 15°C, n = 7 *tbx-35(-)* 25°C. (E) HLH-1::GFP per MS8 sublineage in each *ceh-51(-)* embryo at 20°C. n = 5 *ceh-51(-)*. (F) Best sustained PHA-4::GFP (left) and HLH-1::GFP (right) per MS8 sublineage in *tbx-35*(−) maternal 25°C (light red), *tbx-35*(-); rescuing array (blue, pooled across rearing temperatures), and ceh-51(−) (purple). Boxes are median/IQR, points are per-embryo. PHA-4: n = 9 / 5 / 4; HLH-1: n = 7 / 3 / 5. p values from pairwise Wilcoxon rank-sum, BH-corrected.

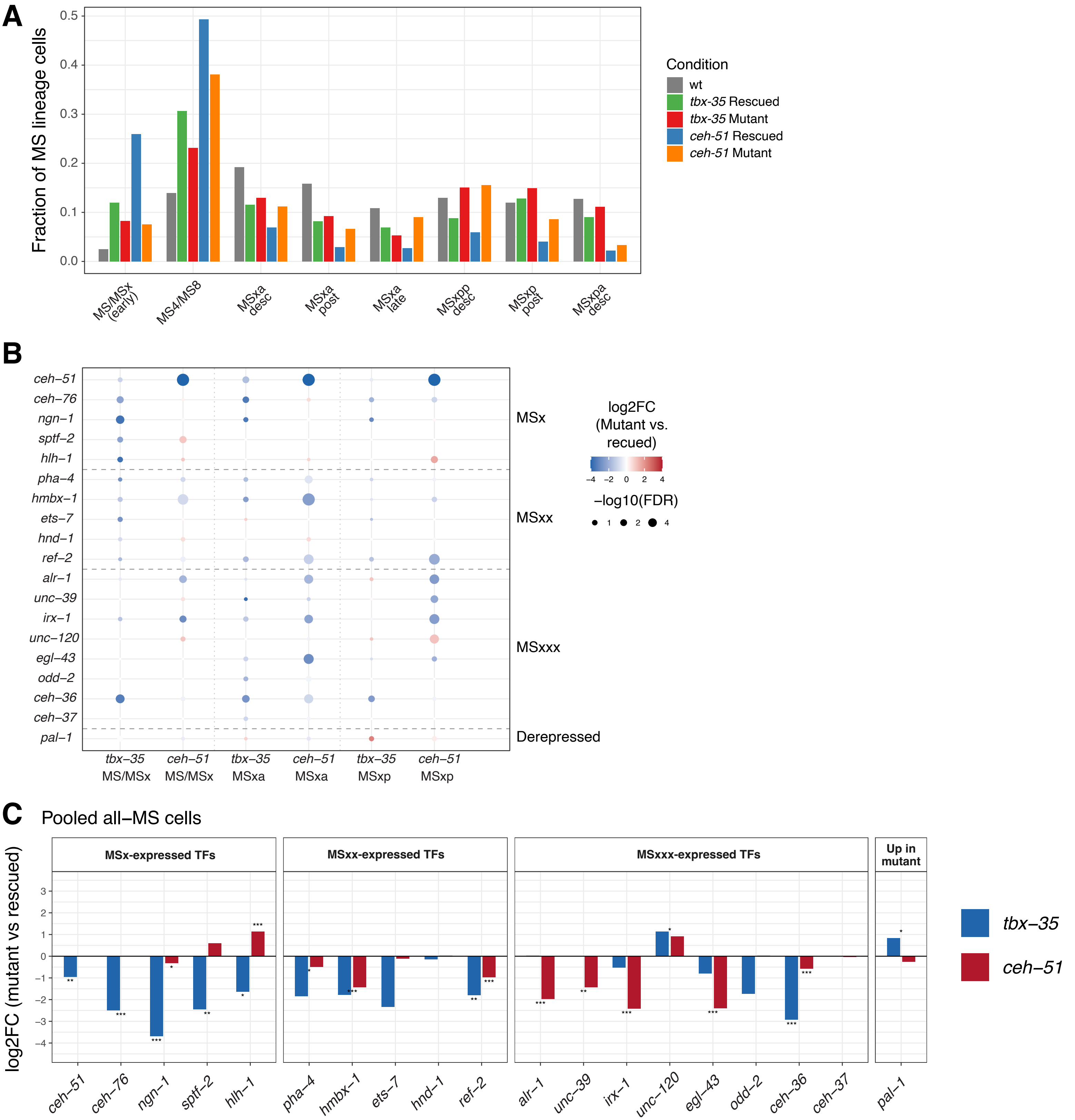


Supplementary Figure 6

**Supplementary Figure 6. Analysis of mutant single-cell data**

(A) Fraction of cells in each MS lineage subcluster derived from each annotation (mutant, rescued, wild-type). (B) Lineage-subdivided expression changes in each mutant. (C) DEG tests for genes initially expressed at each stage in the MS lineage. In these comparisons, DEG are calculated for all cells in the MS lineage clusters.
